# Multi-Modal photoFRESH: Light-Pipe Embedded Printing of Heterogeneous Hydrogel and Tissue Architectures

**DOI:** 10.64898/2026.07.19.738036

**Authors:** Caner Dikyol, William B. O’Brien, Maria A. Stang, Syed Faaz Ashraf, Durva Naik, Jacqueline M. Bliley, Adam W. Feinberg

## Abstract

Recreating the complex spatial gradients and multi-material transitions of native tissues remains a fundamental challenge in 3D bioprinting. To address this, we introduce multi-modal photoFRESH, which integrates localized photochemistry into embedded printing by delivering light through a fiber-optic light-pipe. By tuning numerical aperture, print speed, and photoabsorber content, we achieve precise layer-by-layer control of crosslinking, stiffness, and bioorthogonal biomolecular tethering while preserving high print fidelity. Both photoactivatable support baths and extruded bioinks can be patterned, together with traditional FRESH printing. Utility of the platform is demonstrated by the fabrication of structurally complex tissue scaffolds and cellularized muscle constructs with distinct mechanical and biochemical domains. This multi-modal approach expands the boundaries of embedded bioprinting toward functional and heterogeneous tissue architectures.

## MAIN TEXT

Since the commercialization of stereolithography (SLA) in the 1980’s, 3D printing has rapidly evolved from a rapid prototyping tool and into a widely used advanced manufacturing process across industries. This transition has been enabled by continuous improvements in the materials, robotic control, and processing methods being used, with new fabrication approaches accelerating adoption and expanding the range of applications. In medicine it is now routine to use SLA to produce anatomical models for surgical planning, digital light processing (DLP) to produce dental aligners, selective laser sintering (SLS) to produce surgical guides and skull plates, and electron beam powder bed fusion (EB-PBF) to produce orthopedic metal implants (*1*). These approaches are clinically successful because they use established biomaterials together with patient-specific design to improve the performance of traditional medical devices. However, adapting these approaches for 3D bioprinting to repair, regenerate or replace diseased and damaged tissue has proved quite challenging, due in large part to the range of material, biological, and structural complexity (*2*).

Rebuilding human tissue requires the ability to recreate structure-function relationships that span from the molecular to organ length scales. On the structural side, 3D bioprinting can already replicate the architecture of native tissues by fabricating scaffolds and cellularized constructs with resolution on the order of 10-100 µm (*3*). Light-, droplet-, and extrusion based bioprinting techniques have all shown steady improvement in structural fidelity and cell viability using a range of hydrogels. However, the hierarchical architecture of native tissues is characterized by transitions in mechanical and biochemical cues in a gradient manner, imparting spatiotemporal heterogeneity that is critical to biological function. This includes diverse processes, from growth factor gradients that drive vascularization to mechanical gradients that distribute stress through the myotendinous junction. For example, in extrusion-based systems, the deposited bioink crosslinks uniformly throughout the bioprinting process, so different extruders are typically required to change biomaterial properties (*4*). For droplet-based and inkjet systems, each nozzle ejects the same biomaterial and thus requires arrays for different droplets to approximate gradients (*5*). In contrast, light-based systems such as SLA and DLP excel at spatially patterning photoactivatable biomaterials across both time and space to recapitulate gradients in stiffness, growth factors, and other properties through controlled photochemical reactions (*6*). The disadvantage for light-based systems is the combining of multiple biomaterial and cell types within a single construct, as this requires using multiple bioresin vats or microfluidic systems that add cost and complexity. Thus, while extrusion-, droplet- and light-based techniques are capable of meeting certain biofabrication requirements, none can currently achieve all of them, which fundamentally limits the complexity and functionality that can be attained.

Multi-modal bioprinting has emerged as an effective strategy for integrating the complementary strengths of different biofabrication modalities within a single manufacturing workflow (*7*). Embedded bioprinting has been a key step in this direction, combining extrusion-based printing within an aqueous support bath to allow the environment to control bioink gelation (*8*). Specifically, Freeform Reversible Embedding of Suspended Hydrogels (FRESH) has significantly increased the shape fidelity of bioprinted constructs as the effect of gravity on the soft biological materials is negated while enabling pH, ionic, enzymatic, thermal, chemical and light-based control of crosslinking (*9, 10*). Combining FRESH and related embedded-printing methods with light-based bioprinting methods has already been recognized as a way to fabricate more functional tissue constructs. Embedded extrusion-volumetric printing combines the deposition of cell-laden bioinks within a photocrosslinkable support bath photopatterned with volumetric printing to rapidly generate multicellular and multimaterial tissues with enhanced structural complexity (*11*). Multiple extruders can be used to achieve multi-material volumetric printing of hydrogels, combining gelatin as a sacrificial biomaterial together with hyaluronic acid and synthetic PEG hydrogels (*12*). Software advances have further addressed some of the optical challenges in embedded-volumetric printing, with adaptive, context-aware and closed-loop, image-guided algorithms to improve precision and spatial fidelity (*13*). Extrusion- and light-based printing have also been combined into a single platform to fabricate thermoplastic scaffolds surrounded by patterned hydrogels in proof-of-concept studies (*14*). Collectively, these advances underscore the growing potential of hybrid manufacturing strategies to produce anatomically and functionally more sophisticated tissue constructs. But challenges remain, as volumetric printing is limited to small build volumes due to light penetration into the print chamber, the need for transparent bioinks, and the difficulty in creating spatial gradients due to the way light is delivered. Thus, to date there still has not been an integrated bioprinting approach that combines the full capabilities of embedded printing with precise light-based spatial control comparable to SLA or DLP.

In this study, we introduce photoFRESH 3D Bioprinting, a new technique that integrates photochemistry with extrusion-based embedded bioprinting **(Fig. 1A)**. In this approach, light is delivered from the tip of a light-pipe that contains a fiber-optic cannula, which is rastered within the support bath in a layer-by-layer manner for spatiotemporally controlled photoactivation **(Fig. 1B-C)**. Combining extrusion-based bioprinting with fiber-optic delivered photocrosslinking has been recognized for its potential synergies, but the capabilities have been limited (*15–17*). Lee et. al. enhanced the mechanical stability of cell-laden methacrylated hydrogels by embedding optical fibers within the center of the nozzle to photocrosslink the bioink during printing (*15*). Further, Cianciosi et. al. demonstrated photopatterning with an LED-coupled optical fiber within a photosensitive-laden vat for photocrosslinking hydrogels (*17*). Here, by modulating several parameters of optical-fiber coupled light path (such as rastering speed, light intensity, the wavelength of delivered light) or by changing the formulation of photoactivatable material (such as support bath or extruded bioink), we demonstrate the integration of photochemistry into the embedded platform. We present photocrosslinking of biomaterial inks and support baths, photoconjugation of small molecules, multi-wavelength photoactivation, and multi-modal bioprinting combining light and extrusion. We then apply these capabilities to fabricating tissue engineering scaffolds and cellularized muscle constructs.

**Figure 1.**
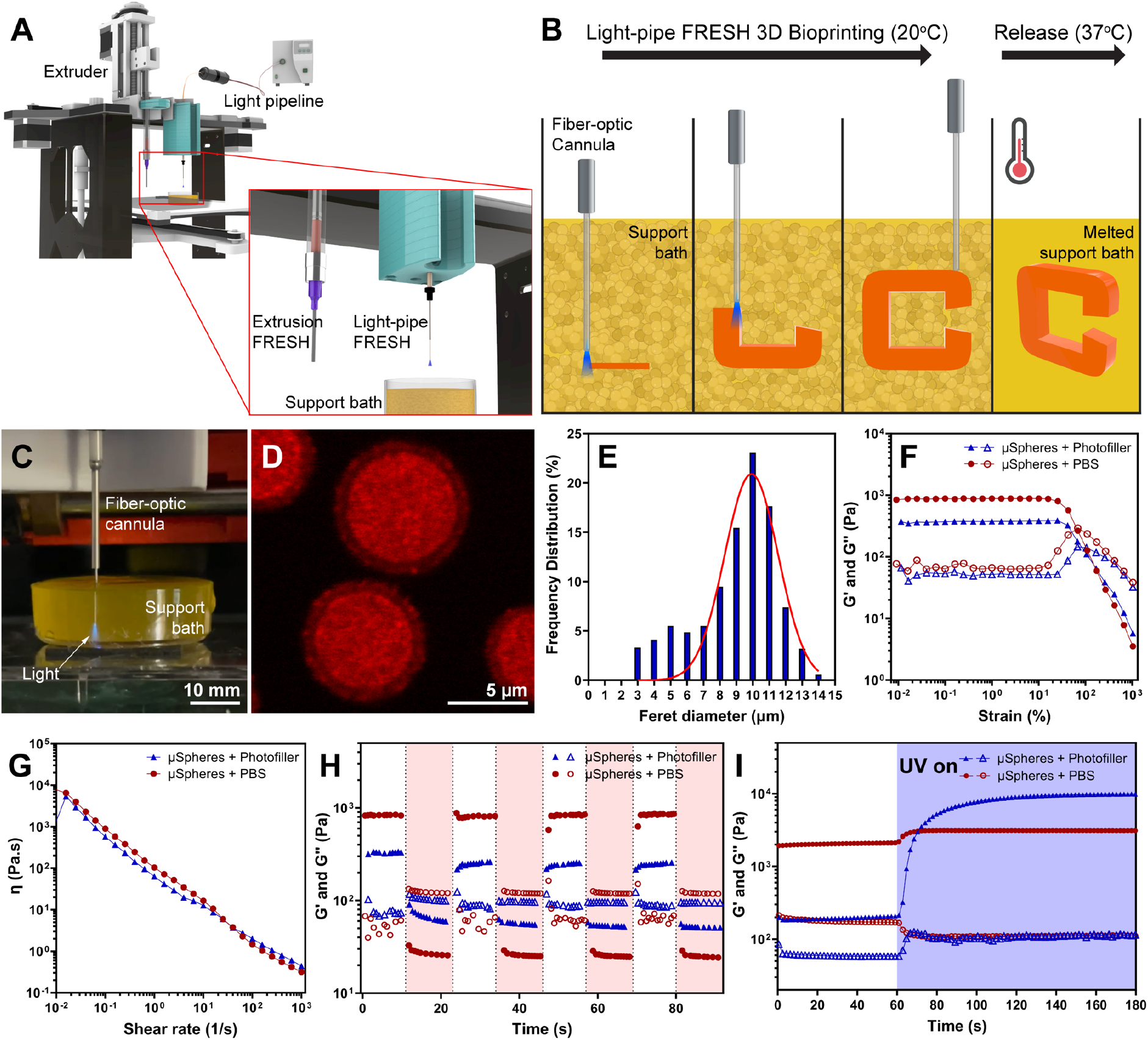
The photoFRESH 3D bioprinting process and gelatin methacroyl (GelMA) support bath characterization. **(A)** Schematic of the photoFRESH 3D bioprinting technique composed of a photoactivatable support bath and a light-pipe integrated into a 3D bioprinter. **(B)** A schematic of the photoFRESH 3D bioprinting process showing the GelMA granular support bath being locally photoactivated (orange) through the light delivered from the tip of a fiber-optic cannula. The 3D object is built layer by layer and, when completed, is released by heating to 37°C and melting the non-photocrosslinked GelMA support bath. **(C)** Image of the photoFRESH setup showing the fiber-optic cannula of the light-pipe placed within the support bath and delivering UV light. (**D**) Confocal image of the GelMA microspheres successfully generated through the complex coacervation process. (**E**) Diameter distribution of the GelMA microspheres. (**F**) Oscillatory amplitude sweep confirms that the support bath maintains similar rheology and yield stress when the photofiller is added. (**G**) Measuring viscosity as a function of shear rate confirms that the shear-thinning behavior of the support bath is also preserved. (**H**) Cyclic strain measurements also confirmed the ability to repeatably yield and recover. (**I**) Photorheology of GelMA microspheres shows that the photofiller improves the crosslinking speed and magnitude. For F-I, storage (G′) modulus is filled symbols and loss (G″) modulus is open symbols.

### Development of a photocrosslinkable microsphere support bath

In photoFRESH, the support bath is composed of photocrosslinkable gelatin microspheres and becomes one of the biomaterials that can be printed. Similar to standard FRESH printing, the support bath is compacted into a slurry that acts as a yield stress material during printing (~22 °C), the difference is that the light-pipe selectively photocrosslinks defined regions of the gelatin support bath to prevent melting upon raising the temperature (>35 °C). To do this, we first synthesized gelatin methacryloyl (GelMA) as the main photoactivatable material of the support bath, a widely used biomaterial with cell binding and enzymatically-cleavable motifs (*18*). Chemical modification of the GelMA was confirmed by proton nuclear magnetic resonance (1H NMR) spectroscopy (**fig. S1**) with a degree of functionalization of 82.45 ± 2.05% by a 2,4,6-trinitrobenzene sulfonic acid assay. Photocrosslinkable microspheres were formed by complex coacervation based on the FRESH v2.0 method (*10*), combines GelMA, unmodified gelatin, and lithium phenyl-2,4,6-trimethylbenzoylphosphinate (LAP) photoinitiator **(Fig. 1D)** with a 9.91 ± 1.63 µm average diameter **(Fig. 1E**). In addition to the GelMA microspheres, the surrounding aqueous phase was filled with what we term the photofiller, which is composed of soluble GelMA, LAP and a tartrazine photoabsorber **(fig. S2)**. The photofiller was found to be critical, as it fills the interstitial space between GelMA microspheres, increasing the photocrosslinking speed and mechanical integrity of the printed part.

The GelMA microsphere support bath demonstrates the yield-stress behavior required for FRESH printing (*10*) and can also be photocrosslinked. Oscillatory amplitude sweeps demonstrated the yield-stress behavior of the support bath, transitioning from an elastic gel state to viscous liquid-like state **(Fig. 1F)**. Further, the shear thinning behavior was maintained with photofiller over a shear rate from 0.01-100 1/s **(Fig. 1G)**. Cyclic strain tests at low and high oscillatory strains demonstrated the stress-yielding, shear-thinning and shear-recoverable characteristics of the GelMA support bath, which are essential for FRESH printing **(Fig. 1H)** (*19*). Finally, using photorheology we showed that the speed of gelation and storage modulus of the GelMA microsphere support bath was significantly faster using the photofiller compared to PBS **(Fig. 1I)**. Based on these results, we selected a GelMA support bath formulation combining GelMA microspheres and photofiller as the photocrosslinkable components to maximize our ability to modulate gelation and fidelity.

### Building the fiber optic-coupled light-pipe

The light-pipe is the second main component of photoFRESH 3D bioprinting. Briefly, a high-intensity light source is collimated and the specific wavelength of light needed for printing is selected using a band-pass optical filter (**Fig. 2A**). A fiber-optic adapter is then used to focus the light into a patch cord that is coupled to a fiber-optic cannula. Numerical aperture (NA) of the fiber-optic cannula determines the angle over which the optical fiber emits light, making it a critical factor impacting the lateral spread, depth, and scattering behavior of light during its travel within the support bath (**Fig. 2B**). We characterized the light penetration in our microsphere-based support bath based on the half-angle, lateral spread, and in-depth attenuation at multiple locations from the light-pipe tip (**Fig. 2C**). Measurements taken at 500 µm **(Fig. 2D)** and 1000 µm **(Fig. 2E)** away from the tip of the fiber-optic cannula demonstrated that using NA 0.1 had the lowest lateral spread. Calculating the mean half-angle confirmed that light delivered from fiber-optic cannulas with smaller NA had lower lateral spread of the light beam **(Fig. 2F)**. Conversely, while light from the fiber-optic cannula with higher NA penetrates less, it exhibits a wider divergence angle compared to its counterparts, delivering more light but also reducing XY resolution. Z-axis light attenuation in the support bath was also correlated with the lateral spread and half-angle measurements, with the higher NA conditions showing more rapid decay in intensity **(Fig. 2G)**. Thus, to maximize spatial resolution in the XY plane, the fiber-optic cannula with 0.1 NA, which showed the lowest lateral spread, was selected for the rest of the photoFRESH 3D bioprinting experiments.

**Figure 2.**
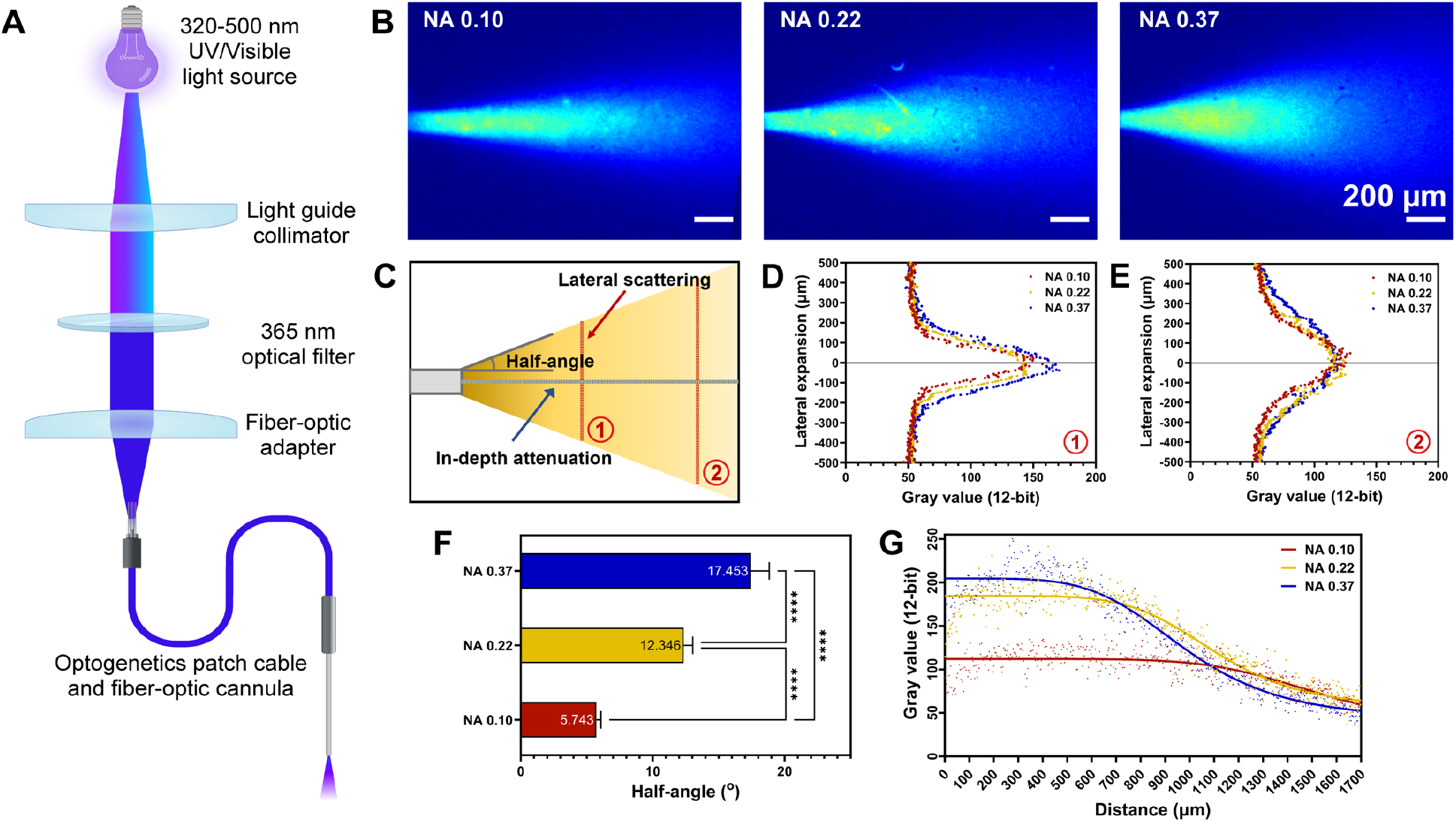
Assembly of components for photoFRESH and optical properties for light delivered from fiber-optic cannulas with different numerical apertures (NA). **(A)** Diagram of the optical path within the light-pipeline setup. **(B)** Fluorescent images of light emitted within the GelMA support bath as a function of NA of the optical fiber in the cannula. **(C)** Schematic of the half-angle, lateral spreading and attenuation measurements. Spread of light laterally at **(D)** 500 µm and **(E)** 1000 µm from the tip of the fiber-optic cannula. **(F)** Half-angle of the different fiber-optic cannulas, showing reduced lateral spread for lower NA optical fibers [N = 3, data are means ± SD, ****P < 0.0001 (One-way ANOVA followed by Tukey’s post-hoc test)]. **(G)** Attenuation of light intensity as a function of penetration depth into the support bath.

### Integration of the Light-Pipe into the 3D Bioprinter for photoFRESH

The fiber-optic coupled light-pipe was integrated into a custom-designed printhead on an open-source 3D printing platform **(fig. S3)**, adapted from our previously published designs (*20, 21*). PhotoFRESH uses the established extrusion-based printing process, where 3D models are designed in CAD and sliced into G-code to control light-pipe motion in 3D space. However, there are no extrusion commands, instead to turn the light on and off, the G-code used to turn the fan for cooling the extruder on and off was repurposed to control the optical shutter from the light source. The amount of light exposure was controlled through changing print speed, dwell time, and power output of the light source. For the majority of our studies, we kept the power output constant and changed print speed because this simplifies the G-code. Changing light exposure through the diameter and numerical aperture of the fiber-optic cannula, the wavelength of light, and other parameters will all impact the printing, a portion of which we demonstrate here. The yellow-colored photoabsorber tartrazine incorporated into the support bath, previously shown to be biocompatible and effective in DLP 3D bioprinting, was used to absorb excess light and confine the photopolymerization process to a defined spatial location (*22*). The end result is a 3D printing process where the light-pipe is translated through the photocrosslinkable support bath and emits light in selective regions to photopolymerize the GelMA (**Movie S1**). Following printing, the support bath is warmed to 37 ºC in order to melt and remove the non-photocrosslinked GelMA.

### Tuning photoabsorber concentration and print process parameters

We first investigated the effect of printing parameters such as speed and photoabsorber concentration on print fidelity and dimensional accuracy. To do this we designed a scaffold for engineering muscle tissue constructs, where we could measure the length and thickness of the overall print as well as the filament width within the infill region (**fig. S4A**). Without any photoabsorber there was poor fidelity, however, incorporating tartrazine in the photofiller of the support bath improved fidelity and confirmed that the molar concentration in the support bath could be increased until lateral filament fusion was lost at 100 µM (**fig. S4B**). To tune print parameters and optimize fidelity further, light power was kept constant while photoabsorber concentration was varied from 0 to 100 µM and the amount of light delivered per unit time was controlled by varying printing speed from 25 to 40 mm/min. Results show that both higher photoabsorber concentration and faster print speeds decreased overcuring but can also lead to filament-to-filament delamination in the extreme. This is because filament width was the most sensitive physical feature to changing the print parameters (**fig. S4C**) and had a comparable effect to changing filament width in extrusion-based 3D printing. Scaffold thickness was also highly dependent (**fig. S4D**), but in this case it was the amount of overcuring in the Z-axis with higher tartrazine amounts reducing this. Finally, scaffold length was the least dependent as expected (**fig. S4E**), because this was primarily a function of the printer XY motion system. For the specific tissue scaffold CAD design used here we selected 60 µM tartrazine in the photofiller to obtain the desired print dimensions (**Movie S2**). However, similar to 3D printing in general, the exact parameters selected for a specific 3D print will be dependent on the CAD design, biomaterials, cells, and other factors.

### Modulating hydrogel crosslinking and mechanical properties

In photoFRESH, modulating the amount of light being delivered defines the size and crosslink density of the structures printed from the GelMA support bath, and both parameters impact the effective mechanical properties. To demonstrate this, a clover shaped construct having four domains was printed with the same light power but at different translation speeds of the light-pipe of 25, 30, 35, and 40 mm/min (referred to as F25, F30, F35, and F40, respectively) **(Fig. 3A)**. We measured the effective elastic modulus of these four domains using a microindentation system to produce force-displacement curves, showing a clear difference in mechanical response **(Fig. 3B)**. As expected, based on the visual differences in filament width, the domain printed the fastest at 40 mm/min was the softest with an effective elastic modulus of ~5 kPa that increased to ~27 kPa for the domain printed at 25 mm/min **(Fig. 3C)**. In addition to varying print speed, changing the power output of the light delivered (from 6 to 10 µW) at constant print speed similarly increased crosslinking and the effective elastic modulus (from ~15 to 50 kPa) **(Fig. 3D)**. These results confirmed that it is the amount of light delivered within a given volume as a function of time that controls the amount of GelMA support bath that is photocrosslinked. Further, we measured the effective elastic modulus which depends on the filament diameter and crosslink density as well as the spacing and geometry of the lattice infill that was indented. Differences at the filament level were confirmed using fluorescent imaging, which showed smaller diameters and lower fluorescence intensities at the faster print speeds indicating less material was photocrosslinked **(fig. S5)**.

**Figure 3.**
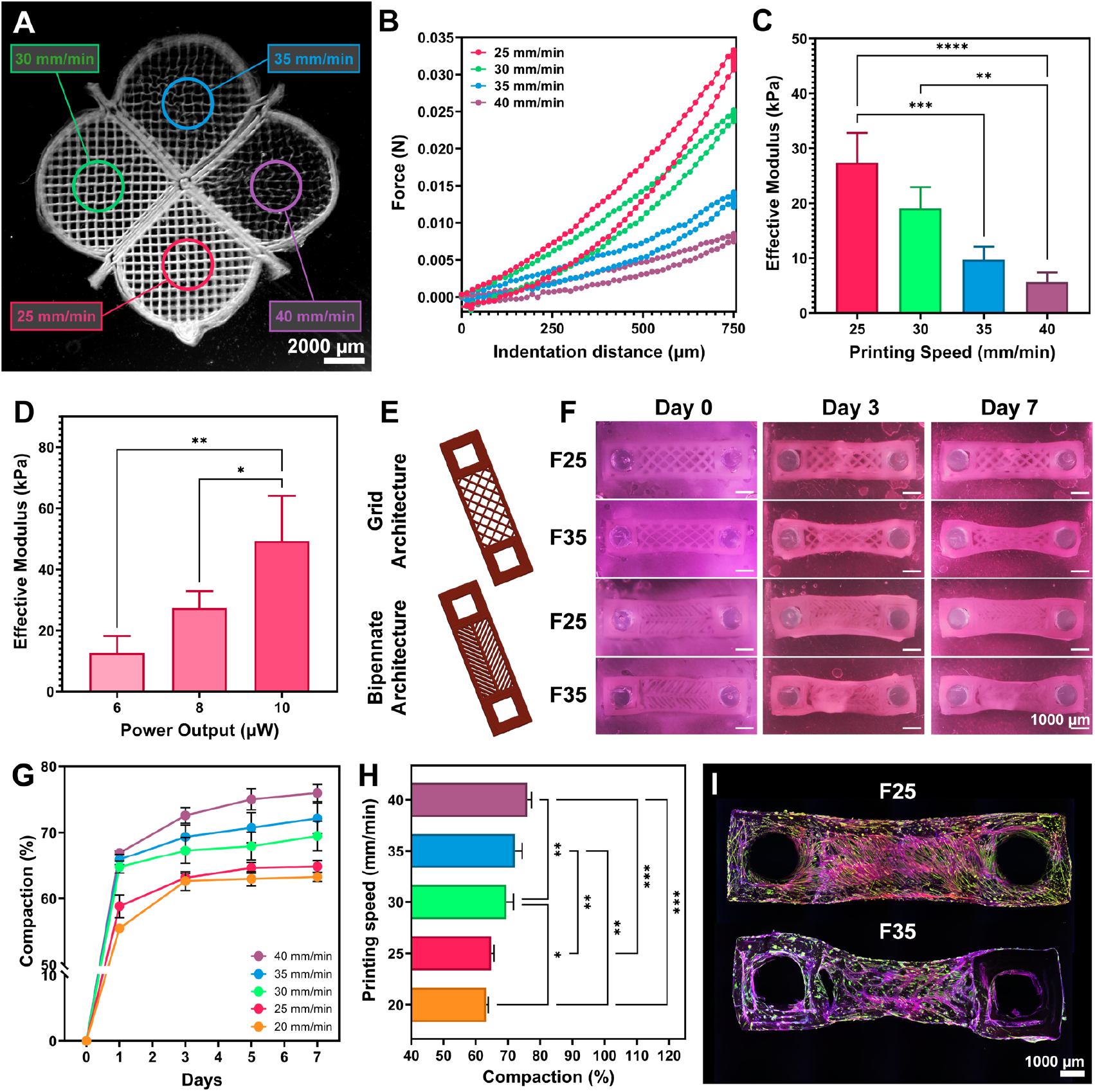
Tuning printing parameters to control mechanics and compaction of skeletal muscle tissue constructs. **(A)** Clover-shaped microindentation sample having four domains bioprinted with constant light intensity at speeds of 25, 30, 35 and 40 mm/min. **(B)** Force versus indentation for each domain of the construct showing that print speed impacts the mechanical properties. **(C)** The effective elastic modulus decreases with increasing print speed when the light power output is held constant [N = 3, data are means ± SD, ****P < 0.0001 (One-way ANOVA followed by Tukey’s post-hoc test)]. **(D)** The effective elastic modulus increases with increasing light power output when the print speed is held constant [N = 3, data are means ± SD, **P < 0.01 (One-way ANOVA followed by Tukey’s post-hoc test)]. **(E)** The g-code rendering of two muscle tissue scaffold designs consisting either a grid or bipennate central scaffold region. **(F)** Time-lapse images at Day 0, 3 and 7 showing cell-mediated compaction of GelMA scaffolds printed at F25 (higher stiffness) or F35 (lower stiffness) and the impact of both print speed and scaffold design. **(G)** Quantified compaction for the grid architecture over 7 days showing that faster print speeds (soft constructs) undergo more compaction. **(H)** Comparison of compaction at Day 7 [N = 3, data are means ± SD, ***P < 0.001 (One-way ANOVA followed by Tukey’s post-hoc test)]. **(I)** Confocal images at Day 19 reveal differences in the way cells grow on and into the scaffolds as a function of printing speed (blue: nuclei, green: myosin heavy chain, red: F-actin).

Next, we investigated how cells responded to these changes in scaffold mechanical properties, with either a grid architecture infill (90º angle square lattice) or a bipennate architecture infill (±45º parallel filaments) **(Fig. 3E)**. Scaffolds were fabricated at print speeds from 20 mm/min to 40 mm/min to tailor the mechanical properties and then placed within PDMS molds with two posts. Scaffolds were then seeded with mouse C2C12 myoblasts in a collagen gel, within which the cells generated uniaxial tension during cell mediated remodeling. Over 7 days of active compaction followed by differentiation up to Day 19, the skeletal muscle cells compacted around and infiltrated the GelMA scaffolds, with changes in shape based on both print speed and scaffold design **(Fig. 3F)**. The amount of compaction ranged from ~60% to ~75%, depending on the effective modulus of the scaffold, **(Fig. 3G), (Fig. 3H)**. Immunofluorescent staining revealed that skeletal muscle cells remodeled the softer scaffolds differently than the stiffer ones, appearing to migrate and grow deeper into the softer construct **(Fig. 3I, fig. S6)**. Functionally, live calcium imaging confirmed that both scaffolds produced constructs with electrically active myotubes that could be stimulated to contract **(Movies S3 and S4)**.

### Spatial patterning of scaffold mechanics influences contractility of engineered muscle tissue

Using photoFRESH, we next modulated mechanical properties within scaffolds, comparing cell response to stiff/stiff (F20-F20), soft/stiff (F35-F20), and soft/soft (F35-F35), conditions. There was more compaction of the softer F35 domains compared to the stiffer F20 domains, with the F35-F20 scaffold showing a clear transition at the interface **(Fig. 4A)**. Clear differences in cell infiltration were also observed, with cells migrating into and aligning on the lattice infill filaments of the F35 scaffold as compared to staying on the surface and not aligning to the filaments of the F20 scaffold. While the muscle constructs all had the same length when in the PDMS molds, removing the constructs from the molds revealed that the softer F35 domains deformed further due to cell-generated pre-stress while the stiffer F20 domains maintained dimensions **(Fig. 4B)**. To better understand why these differences occurred, we recorded the calcium transients in the cells under field stimulation across the soft and stiff domains showing clear variations **(Fig. 4C)**. Focusing in on a single stimulation, the calcium cycled into the cells increased as the scaffold became softer, with the F35-F20 scaffold have intermediate values **(Fig. 4D)**. Quantifying this confirmed a statistically significant increase in the peak calcium levels cycling into the cell during each contraction **(Fig. 4E)**. Because calcium levels are associated with contractility in skeletal muscle cells, we next assessed active deformation during contraction. At baseline, the length of each scaffold type was different, and showed that the softer condition (F35) shortened by >15% due to the pre-stress generated by the cells, while the stiffer condition (F20) shortened by <3% (**Fig. 4F**). Under externally applied 1Hz field stimulation, the soft F35 regions of the scaffolds visibly shortened during contraction while the stiffer F20 regions did not (**Fig. 4G** and supporting **Movie S5**). Quantifying this deformation confirmed that the softer scaffold (F35-F35) contracted the most, the stiffer scaffold (F20-F20) contracted the least, and the soft/stiff scaffold (F35-F20) was intermediate **(Fig. 4H)**. Considering the calcium and contraction data together suggests that at least in part, the increased contractility is due to an increase in peak calcium cycling as a function of scaffold mechanical properties, in line with previous findings (*23, 24*). These results demonstrate how the mechanical gradients generated by light-pipe FRESH 3D bioprinting can affect the pre-stress, contractile shortening and calcium cycling behavior of skeletal muscle cells. It also highlights the ability of photoFRESH to modulate scaffold mechanical properties by simply changing the amount of light delivered during photocrosslinking with spatial control, something that is not possible with existing extrusion-based embedding printing methods.

**Figure 4.**
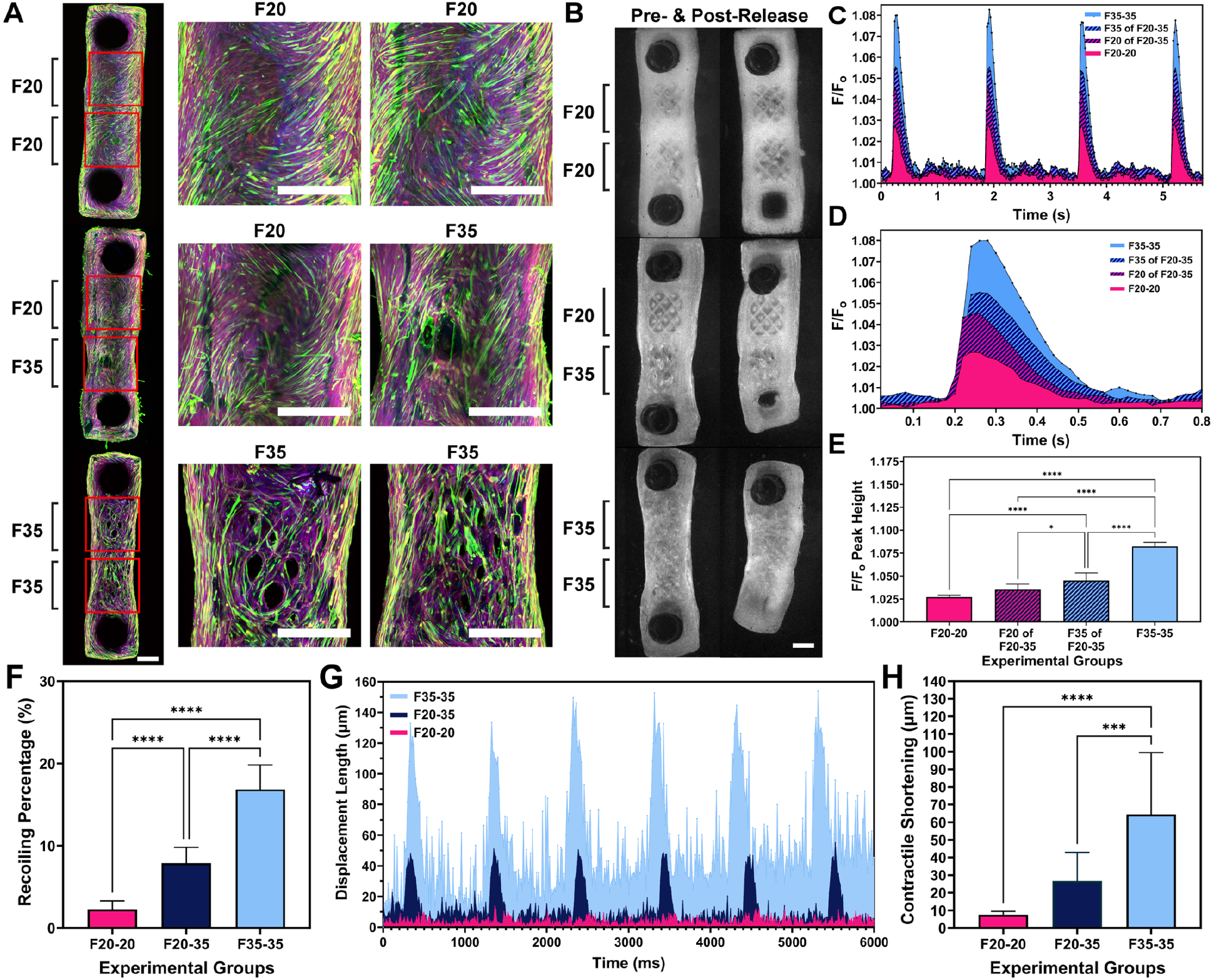
Scaffold mechanics affect cell remodeling, calcium handling and contractility of muscle tissue constructs. **(A)** Confocal imaging of constructs with stiff/stiff (F20/F20), stiff/soft (F20/F35) and soft/soft (F35/F35) domains shows that cells preferentially migrate onto the GelMA filaments within the soft (F35) regions (blue: nuclei, green: myosin heavy chain, red: F-actin) (scale bars: 1 mm). **(B)** Releasing the constructs from the one of the posts shows distinct difference in the elastic recoil due to cell-generated pre-stress in the scaffolds (scale bars: 1 mm). **(C)** Contractions were paced under 1 Hz field stimulation, and calcium imaging was used to measure calcium cycling in and out of the cells. **(D)** Focusing in on a single contraction, comparison of relative fluorescence at 1Hz stimulation shows that the magnitude of calcium cycling into the cells depends on scaffold print speed. **(E)** The peak calcium when field stimulated to twitch [N = 3, data are means ± SD, ****P < 0.0001 (One-way ANOVA followed by Tukey’s post-hoc test)]. **(F)** The magnitude of elastic recoil increases with print speed, likely due in part to scaffold stiffness [N = 3, data are means ± SD, ****P < 0.0001 (One-way ANOVA followed by Tukey’s post-hoc test)]. **(G)** Field stimulation of the muscle constructs at 1 Hz produces larger deformations in the softer scaffolds. **(H)** The magnitude of contractile shortening when field stimulated to twitch confirms this [N = 3, data are means ± SD, ****P < 0.0001 (One-way ANOVA followed by Tukey’s post-hoc test)].

### Multi-modal light-pipe combined with extrusion bioprinting to build complex scaffolds

To achieve true multi-modal 3D bioprinting capabilities, we next combined photoFRESH with extrusion FRESH within the same support bath for multi-material biofabrication of complex tissue scaffolds. This enables both the support bath and the extruded bioinks to be part of the scaffold structure, and to use photochemistry to spatially pattern distinct mechanical and biochemical domains within these biomaterials. To demonstrate this, we first designed a multi-pennate muscle-inspired architecture with a myotendinous junction-like interface between tendon (extruded collagen) and muscle (light-pipe GelMA) regions **(Fig. 5A)**. As noted in the schematic, this design combines two entirely different printing modalities (FRESH and photoFRESH) to produce a single integrated construct with multiple biomaterials, architectures, and mechanical properties. The extruded collagen was not chemically-modified and not photocrosslinkable, meaning it gels through self-assembly into fibrils during FRESH printing as we have previously demonstrated (*10, 25*), thus, we designed the scaffold to have a mechanical interlock for integration with the light-pipe printed GelMA. Results confirmed the ability to print the tendon-like domain of aligned collagen filaments interfaced directly with the muscle-inspired domains of light-pipe printed GelMA support bath **(Fig. 5B, Movie S6)**. Zooming in on the interface between domains showed the ability to precisely control biomaterial placement with each printing method, and to maintain structural integrity of the entire construct, similar to the interlocking between jigsaw pieces.

**Fig. 5.**
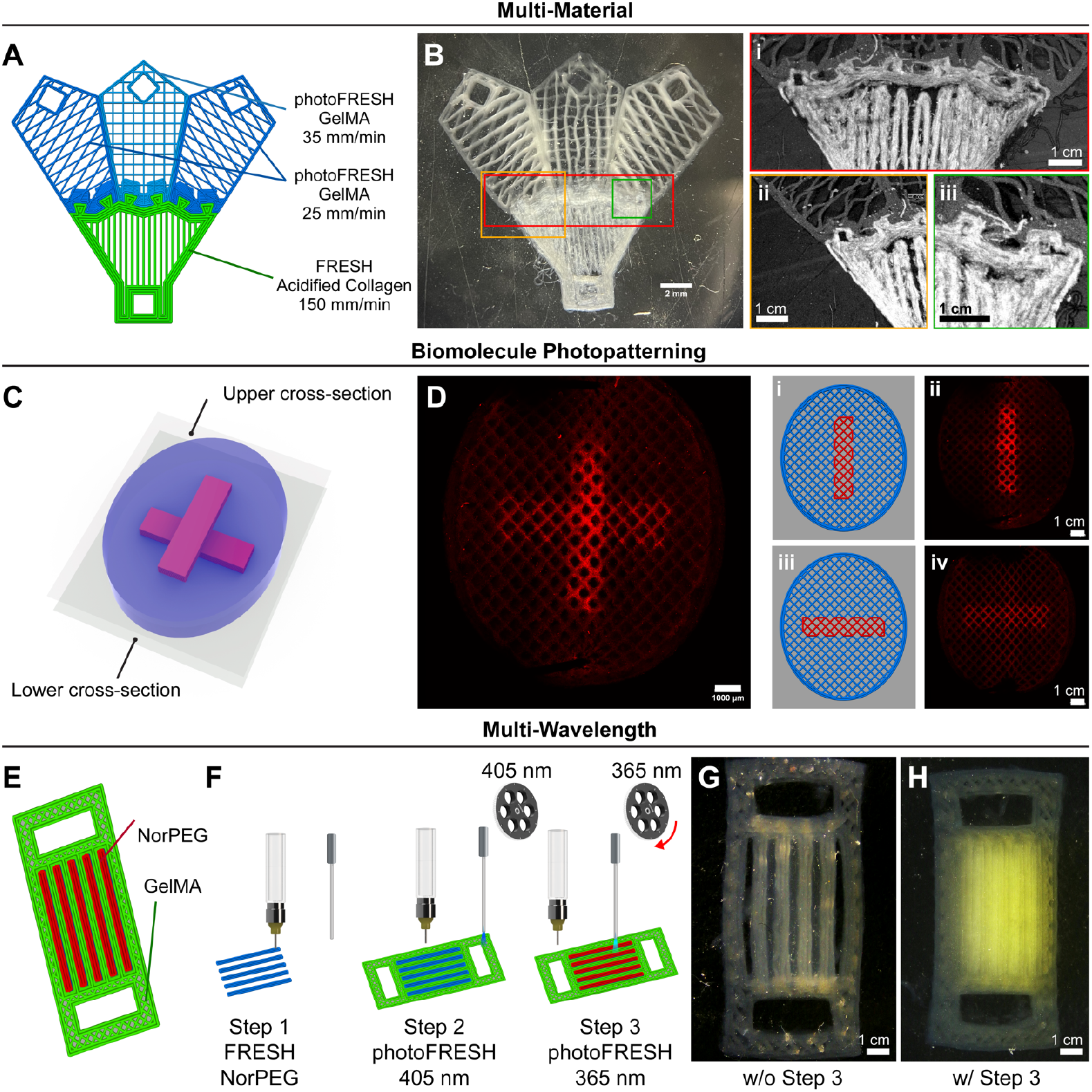
Multi-modal printing combining photoFRESH and FRESH in the same support bath for multi-material and multi-wavelength biofabrication. **(A)** Schematic of myotendinous junction inspired scaffold architecture with the dual material tendon-like domain consisting of extruded collagen type I and the muscle-like domains consisting of light-pipe exposed GelMA. **(B)** Stereomicrograph of multi-material bioprinted scaffold with **(i-iii)** closer view of the interface between the FRESH printed collagen and photoFRESH printed GelMA imaged by OCT. **(C)** Schematic of a scaffold designed as a FRESH printed collagen disc with two domains photopatterned onto the collagen in different Z-planes with photoFRESH. **(D)** Top-down max intensity confocal image of the fabricated scaffold showing the two domains of methacrylated rhodamine photopatterned within the methacrylated collagen scaffold, with **(i-iv)** images of the specific Z-planes within the scaffold confirming axial control. **(E)** Scaffold design consisting of alternating domains of photocrosslinked GelMA support bath along with extruded and photocrosslinked NorPEG. **(F)** Multi-wavelength bioprinting process. Step 1, NorPEG was extruded into the GelMA support bath. Step 2, GelMA support bath was photocrosslinked with visible light coming from the light-pipe. Step 3, extruded NorPEG was photocrosslinked with the UV light delivered from the light-pipe after filter switches. **(G)** Stereomicroscope image of the bioprinted scaffold without Step 3, showing that the NorPEG will dissolve if not photocrosslinked. **(H)** Stereomicroscope image of the bioprinted scaffold with Step 3, confirming that the NorPEG is stabilized when photocrosslinked.

Next, we used photoFRESH not to print the GelMA support bath but instead combined it with FRESH to spatially pattern biomolecules in 3D onto a collagen scaffold. To demonstrate this, we used the standard FRESH gelatin support bath (not GelMA) and added methacrylated rhodamine into the solution phase. With this approach, we FRESH printed methacrylated collagen bioink and then used photoFRESH to spatially control crosslinking of methacrylated rhodamine. The test scaffold consisted of a FRESH printed collagen disk with two photoFRESH patterned orthogonal rectangular regions of rhodamine separated in the Z-axis **(Fig. 5C)** and was printed by alternating between the collagen extrusion and light-pipe print heads on the printer, layer-by-layer **(Movie S7)**. Confocal imaging of the printed scaffold confirmed that light delivered from the fiber-optic cannula on specific locations of the extruded collagen filaments was effective at photopatterning within the XY planes and between Z planes **(Fig. 5D)**. This establishes that photoFRESH can be used beyond crosslinking hydrogels and support baths, and that it is also capable of photoconjugation for spatial patterning onto FRESH printed scaffolds.

Finally, to establish the full multi-modal printing capabilities of photoFRESH, we combined FRESH with photoFRESH using bio-orthogonal photoactivation wavelengths and photochemistries all applied together within the same construct. To perform multi-wavelength photoFRESH, we added a motorized filter wheel to the light-pipe bioprinting system with the ability to switch between up to 6 filters, and here used 365 nm and 405 nm band-pass filters for proof-of-concept **(fig. S7)**. We designed a construct consisting of alternating regions of photoFRESH printed GelMA support bath along with FRESH printed norbornene-functionalized PEG (NorPEG) that was also photocrosslinkable (**Fig. 5E)**. The frame of the construct was photoFRESH printed GelMA support bath photocrosslinked at 405 nm in the presence of Ru/SPS photoinitiator in the support bath **(Fig. 5F)**. NorPEG was FRESH printed in between the main struts of the GelMA scaffold. The extruded NorPEG was then photocrosslinked using photoFRESH at 365 nm in the presence LAP photoinitiator in the support bath. As a control experiment, if the second photoFRESH step to photocrosslink the NorPEG was not performed, then the FRESH printed NorPEG diffused away while melting the GelMA support bath during the release process (**Fig. 5G**). In comparison, adding photoFRESH with UV light at 365 nm in the presence LAP photoinitiator enabled photocrosslinking of the extruded NorPEG through the secondary wavelength delivered from the light-pipe (**Fig. 5H**). These results confirm that photoFRESH can be used to photocrosslink the support bath, photocrosslink FRESH printed bioinks, and photopattern onto FRESH printed bioinks.

In summary, photoFRESH 3D bioprinting integrates precise spatial control of photochemistry within embedded 3D printing. Through fiber-optic delivery of light into the support bath or extruded bioink via a light-pipe, this technique enables patterning of bioinks, scaffold mechanics and biomolecules with a resolution on the order of ~100 µm while maintaining all of the advantages of embedded 3D bioprinting. These results demonstrate that within the same support bath, we can combine FRESH extrusion-based and photoFRESH multi-wavelength light-pipe printing in an integrated process, greatly expanding both the range of biomaterials and spatial patterning that can be achieved in a biofabrication system. While we have used muscle-inspired scaffold designs for proof-of-concept this technique is broadly adaptable for advanced biofabrication applications, providing a new methodology for replicating the spatial structural and functional heterogeneity of human tissues.

## Author contributions

Conceptualization: C.D., A.W.F. Methodology: C.D., W.O., M.A.S., S.F.A., D.N., and J.M.B Investigation: C.D., W.O., M.A.S., S.F.A., D.N., and J.M.B. Visualization: C.D., W.O., and S.F.A. Funding acquisition: A.W.F. Project administration: A.W.F. Supervision: A.W.F. Writing – original draft: C.D. and A.W.F. Writing – review & editing: C.D., W.O., M.A.S., S.F.A., D.N., J.M.B., and A.W.F.

## Data and materials availability

All data needed to evaluate the conclusions in the paper are present in the main text and the supplementary materials. The STL files for 3D bioprinter hardware modifications and printed constructs are available under an open-source CC-BY-SA license.

## Supplementary Materials

Materials and Methods

Figs. S1 to S9

Movies S1 to S7

