## Supplemental Information for "Multi-Modal photoFRESH: Light-Pipe Embedded Printing of Heterogeneous Hydrogel and Tissue Architectures"

**This PDF file includes:**

Materials and Methods
Supplemental Information References
Figs. S1 to S9
Captions for movies S1 to S7

**Other supplementary material for this manuscript includes:**

Movies S1 to S7

### MATERIALS AND METHODS

#### Gelatin Methacryloyl Synthesis and Characterization

Gelatin methacryloyl (GelMA) was synthesized through the reaction of lysine and hydroxylysine groups of gelatin with methacrylic anhydride (MA), based on previously described methods (*1*). Briefly, 10% (w/v) gelatin type B (Sigma Aldrich) was dissolved in phosphate buffered saline (PBS) (Sigma Aldrich) at 60 ºC and stirred until the solution became transparent. MA (Sigma Aldrich) was added dropwise into dissolved gelatin solution in the chemical hood, at a ratio of 0.6 g of MA per 1 g of gelatin. The solution was stirred for 3 hours using an overhead stirrer and then centrifuged at 3500 rpm for 3 minutes to remove unreacted MA. The GelMA in the supernatant solution was diluted 1:1 in PBS at 40 ºC and then dialyzed using 12 kDa membranes in ultra-pure Milli-Q water for one week. Finally, the GelMA solution was frozen at -80 ºC overnight and then lyophilized for 3 days prior to storage at -20 ºC until further use. Successful functionalization of gelatin with MA was characterized by 1H NMR using a Bruker Avance II spectrometer **(fig. S1)** by dissolving gelatin and GelMA in deuterium oxide, $D_2O$ (50 mg/mL) at 40 ºC and analyzed using Bruker TopSpin software. The degree of functionalization (DoF) for the synthesized GelMA was characterized based on the 2,4,6-Trinitrobenzene Sulfonic Acid (TNBS) assay (*2*). Briefly, gelatin and lyophilized GelMA samples were dissolved within 4% (w/v) sodium bicarbonate (Sigma Aldrich) at 1 mg/mL concentration. Then, 0.5 mL of 0.2% (w/v) TNBS (Sigma Aldrich) was added to 0.5 mL of the dissolved gelatin and GelMA. After rigorous mixing, the solutions were allowed to react at 37 ºC for 2 hr. The reaction was quenched by adding 0.5 mL of 10% (w/v) sodium dodecyl sulfate (Sigma Aldrich) and 0.25 mL of 1M HCl. The absorbance of the samples was then measured at 335 nm using a plate reader. The molar concentration of free primary amino groups was compared to glycine standard solutions at concentrations of 0, 2, 4, 8, 16, 32, and 64 µg/mL. DoF was calculated as $(A_0 - A)/A0 \times 100\%$, where $A_0$ and A were the absorbance values of the native and functionalized gelatin, respectively.

#### Generation of Photocrosslinkable Granular Support Bath

The photocrosslinkable support bath was made out of GelMA-based microspheres surrounded by a photofiller composed of GelMA, photoinitiator and photoabsorber. Photocrosslinkable microspheres were generated by modifying the previously published coacervation process to create gelatin microspheres (*3*), with GelMA added as a source of photoreactive groups and lithium phenyl-2,4,6-trimethylbenzoylphosphinate (LAP) as a photoinitiator. Briefly, 0.1% (w/v) of LAP was dissolved in distilled water at 70 ºC for 30 min, and an equal volume of 200 proof ethanol was then added. Next, 1.5% gelatin type B (Sigma Aldrich), 5% GelMA and 0.25% Pluronic F127 (Sigma Aldrich) were added and dissolved, followed by the addition of 0.6% gum Arabic (Sigma Aldrich). The pH was then adjusted to 5.5 using NaOH, stirred overnight, and then held at room temperature for 6 hours to allow the microspheres to settle out of solution. The liquid phase was removed and the microparticles were washed with distilled water and 1X PBS. The aqueous phase surrounding the microspheres was then replaced with a photocrosslinkable solution made out of 9% GelMA, 0.5% LAP and varying concentrations of tartrazine, which we refer to as the photofiller. To do that, photofiller was mixed with an equal volume of compacted microspheres, centrifuged at 4500 g for 10 min, and then the supernatant was removed. The final compacted GelMA microparticle slurry was transferred into a dish to be used as a photocrosslinkable support bath.

### Rheological Characterization of Support Bath Composition

The viscoelastic behavior of the photocrosslinkable support bath was characterized using a DHR-2 rheometer (TA Instruments) equipped with a 20 mm diameter parallel plate geometry and a Peltier plate for temperature control. For each rheological measurement, top geometry was lowered to 500 µm gap distance after the sample was loaded onto the lower geometry and then the contact of samples with air was prevented by covering geometry gap with a low viscosity silicone oil. Linear viscoelastic (LVE) region of the support bath samples was determined by strain sweep from 0.01-100% at 10 rad/s. A 1% strain was employed for the rest of the measurements as it remained within the LVE region for all measurements at different frequencies. The shear thinning behavior of the support bath was also investigated by monitoring changes in the viscosity depending on the changes in shear rate from 0.01-100 1/s. To investigate the support bath's shear recoverability, cyclic strain test at low and high oscillatory strains of 1% and 50% was performed at 10 rad/s angular frequency for 10 seconds per 10 cycles. Photocrosslinkability of the support bath formulation was characterized through photorheology. After 60 seconds of steady measurement of storage and loss modulus at 1% strain and 10 rad/s frequency, UV light was delivered through the glass lower geometry and the evolution of storage modulus over the time was recorded. Results were analyzed using TRIOS (TA Instruments) software and then plotted at GraphPad Prism.

### Assembly of Optical Fiber-Coupled Light Path (Light-Pipeline)

The light pipeline for photoFRESH was composed of multiple optical elements designed to deliver light to a defined region within the support bath (**fig. S8**). The UV-visible light source (Omnicure S2000, Excelitas) was connected by a liquid light guide to a collimator (Mightex). The collimated light passes through a band-pass optical filter (Edmund Optics) in order to transmit the specific wavelength required for the photoinitiator included in the support bath. After the filter, light was coupled into a fiber-optic patch cord having SMA (SubMiniature version A) and zirconia ferrule connector at the ends (Doric Lens). The ferrule end of the fiber-optic patch cord was coupled into a cannula through a mating sleeve (Doric Lens). The fiber-optic cannula consisted of the multi-mode optical fiber within a 100 µm inner diameter, 30 mm long stainless-steel sheathing.

### Investigation of Light Delivery from Fiber-Optic Cannulas with Different Numerical Apertures

The effect of fiber-optic cannula choice was investigated through positioning fiber-optic cannulas with different NAs (0.10, 0.22 and 0.37) laterally onto one side of a cuvette loaded with microspheres and then capturing the emitted light using a stereomicroscope equipped with Prime 95B camera. To be able to capture the UV light delivered, microspheres were washed with DAPI (D3571, ThermoFisher Scientific) beforehand. After capturing the emitted light, attenuation of light in both parallel and vertical to the direction of light penetration was analyzed in ImageJ and plotted in GraphPad Prism.

### Integration of the Light-Pipeline into the Bioprinter and the photoFRESH Process

The light-pipeline was incorporated into a custom-designed, open source 3D printing platform previously developed by our group (*4*). The light-pipeline was mounted onto the printhead using a mating sleeve with a custom-designed adapter. The shutter of the light source was connected to the 3D printer microcontroller through the fan control in order to turn on and off the shutter by G-code control. The 3D models for printing were generated in Rhinoceros 6 (Robert McNeel & Associates, USA), exported as stereolithography (SLA) files and then imported into PrusaSlicer (Prusa3D, USA) slicing software to generate the G-code file. Unless otherwise stated, scaffolds were printed with 365 nm UV light with 8 μW power output using a 100 μm inner diameter fiber-optic cannula and support bath composed of GelMA microspheres photofiller of 9% GelMA, 0.5% LAP and 60 μM tartrazine. After the bioprinting process, the print dish was transferred to a 37 ºC incubator, where the non-photocrosslinked GelMA melted away and printed scaffolds were released. Scaffolds were then rinsed 3 times with 37 ºC 1X PBS supplemented with 1% (v/v) Penicillin-Streptomycin (Pen/Strep).

**Measurement of Effective Modulus by a Custom-Design Microindentation Setup**

The microindentation system consisted of 3-axis horizontal XY and vertical Z motorized stages combined with a force transducer and an indentation rod. The force transducer (ATI Nano17 Titanium) had a load threshold of 12 N in X and Y-axis, and 17 N in the Z-axis. The force transducer was attached to a 1.5 mm diameter cylindrical metal rod with a hemi-sphere fused silica with 1.5 mm diameter (Edmund Optics) at the tip. Force displacement curves were captured by positioning the indenter tip on the surface of the scaffold and moving down 750 μm at 0.1 mm/s while recording from the force transducer via ATI DAQ F/T software (ATI Industrial Automation). The effective elastic modulus of each scaffold domain was calculated from the force-displacement data obtained during indentation using the Oliver Pharr method (*5*, *6*). The maximum force ($F_{max}$), the permanent depth of penetration ($h_f$), and the slope of the upper portion of the unloading curve ($S$) were extracted from the graph. The value for $h_f$ was taken to be the depth in the unloading curve where the force returned to baseline. The value for $S$ was calculated using a linear regression of the upper portion of the unloading curve using MATLAB (MathWorks). The indentation depth ($h_{max}$) and the radius of the indenter ($R$) were both 0.75 mm.

The contact depth, $h_c$. was calculated according to the contact depth equation for spherical indentation:

$$h_c = \frac{h_{max} + h_f}{2}$$

Using this quantity, the microindentation area, $A$, was calculated by:

$$A = 2\pi \mathrm{R} h_c$$

$E_r$, the reduced modulus, was then calculated according to the equation:

$$E_r = \frac{\sqrt{\pi}}{2} * \frac{\mathrm{S}}{\sqrt{A}}$$

In order to determine the effective modulus ($E$), the Poisson's ratio ($v_i = 0.16$) and elastic modulus ($E_i$ = 73 GPa) of the hemi-sphere indenter are taken from the manufacturer datasheet (Edmund Optics, #67-392). Poisson's ratio of the material of interest (GelMA), $v = 0.44$, was taken from the literature (*7*). Then, the $E$ value was calculated according to the following equation:

$$\frac{1}{E_r} = \frac{(1 - v^2)}{E} + \frac{(1 - v_i^2)}{E_i}$$

**Imaging of Filament Surface Morphology**

Confocal images of the filaments from each domain of the clover-shaped construct were taken using a 16X water immersion objective and 4X optical zoom on a Nikon A1R MP+ multiphoton and confocal microscope. Images were captured using GelMA autofluorescence at 555 nm and post processed in ImageJ.

### C2C12 Mouse Myoblast Casting onto Bioprinted Scaffolds

Skeletal muscle tissue casting procedure onto the bioprinted scaffolds was adapted from previously published work (*8*). Briefly, custom-designed PDMS molds were created to house photoFRESH bioprinted scaffolds during casting and culture. Each mold was designed to have a rectangular well with two posts that served as anchor points for the tissue during the compaction period. To create PDMS molds, a negative master-mold was designed and then printed using a plastic 3D printer. PDMS prepolymer mixture was cast into the negative master-molds and then left for curing overnight at 60 ºC. After the curing period, PDMS molds were taken out from the negative master-molds and cleaned by sonication with 70% ethanol. PDMS molds were mounted in 12-well plates with a vacuum grease and then blocked with 1% Pluronic F127 solution. After sterilizing with 15 min UV-ozone treatment, photoFRESH bioprinted scaffolds were transferred to the PDMS molds, where they were anchored around the two posts. C2C12 mouse myoblasts were cultured in growth media (DMEM supplemented with 10% FBS, 1% L-glutamine and 1% Pen/Strep). C2C12 cells were then cast around the tissue scaffolds inside the PDMS molds at a \ concentration of 30 million cells/mL in a solution of neutralized collagen (0.5 mg/mL) and Matrigel (10%). Scaffolds were then incubated at 37°C for 60 minutes to gel the collagen, after which 3 mL of warm growth media was added per well. Media was changed the day after casting and then every other day. On the fifth day after seeding, growth medium was switched to differentiation media (DMEM supplemented with 2% horse serum, 1% L-glutamine and 1% Pen/Strep) in order to direct C2C12 cells towards myotube formation. Stereomicroscope images of the cast tissues were taken during the media change days in order observe the evolution of tissue compaction over the days in between scaffold groups.

### Immunofluorescence Staining and Confocal Imaging

After culturing for up to 21 days, tissue scaffolds were fixed with 4% formaldehyde (15710, Electron Microscopy Sciences) and stained with DAPI (D3571, ThermoFisher Scientific), phalloidin 555 (A22284, Life Technologies) and mouse anti-myosin heavy chain (MA5-11748, ThermoFisher Scientific) primary antibody followed by goat anti-mouse secondary antibody conjugated to AlexFluor488 (A28175, Life Technologies). A Nikon A1R MP+ multiphoton confocal microscope and 4X objective were used to obtain tile scans and 3D z-stacks of all tissues.

### Calcium Imaging of Tissue Scaffolds

Electrophysiology of skeletal muscle tissues was assessed via calcium imaging during spontaneous and stimulated muscle contractions. Tissue scaffolds were incubated in Tyrode's solution with 5 µM calcium indicator Cal 520 AM (21130, AAT Bioquest) and 0.25% Pluronic F-127 (P2443, Sigma Aldrich) for 60 minutes at 37 ºC, followed by a 60-minute incubation at room temperature. After rinsing with Tyrode's solution, tissues were housed within a heated dish at 37 ºC while still being attached to their PDMS chamber. An imaging platform consisting of an epifluorescent stereomicroscope and Prime 95B Scientific CMOS camera was used for high-speed imaging of calcium transients at a frame rate of 80 frames per second. Field stimulation was achieved by placing two parallel platinum electrodes into the dish and using a Grass Stimulator to apply a square wave pulse at a range of frequencies from 1 to 10 Hz (1, 5 and 10 Hz) and 80 V for 20 ms.

### Contractility Measurements

Contractile behavior of tissue scaffolds was analyzed by tracking the displacement of scaffolds under 1 Hz field stimulation. First, one of the posts was removed from the PDMS molds housing the tissue scaffold, allowing the tissue to contract unstrained. Next, the PDMS mold was placed into 37 ºC temperature controlled dish and stimulated with 1Hz field stimulation. The contractile displacement of the tissue scaffold was captured using a stereomicroscope and Prime 95B Scientific CMOS at a frame rate of 80 frames per second. Videos were analyzed to track displacement using IMARIS software and then plotted in GraphPad.

### Dual-Head Bioprinting of Myotendinous Junction-Mimicking Construct

The 3D model of the myotendinous junction-inspired construct was generated in Rhinoceros 6 and sliced in Simplify 3D. In the slicing software, the FRESH extruder was assigned as the primary printhead, the photoFRESH light-pipe was assigned as the secondary printhead. The tool-exchange script was modified to turn on/off the shutter in between every extrusion and photopatterning step. A 35 mg/mL acidified collagen type-1 (LifeInk 240, Advanced Biomatrix) was loaded into a gas-tight glass syringe (1750TLLX , Hamilton) and then mounted onto the 3D bioprinter. A 150 µm inner diameter needle (Jensen Global) was fitted to the syringe and primed. The light-pipe mounted to the bioprinter as the secondary printhead had a fiber-optic cannula with 0.1NA and 100 µm inner diameter (**fig. S9**). To ensure gelation of the FRESH printed collagen bioink, 200 mM HEPES was added to the photofiller rather than 1X PBS. Following the dual-head 3D bioprinting process, the slurry was washed two times with 200 mM HEPES with 1% Pen/Strep. Stereomicroscope and OCT images of the bioprinted construct were taken.

### Photopatterning of Extruded Bioink with Light-Pipe

A photosensitive bioink consisting of 8 mg/mL methacrylated collagen and 11.66 mg/mL acidified collagen was prepared through mixing 12 mg/mL PhotoCol (Advanced Biomatrix) and 35 mg/mL acidified collagen type-1 (LifeInk 240, Advanced Biomatrix). The prepared bioink mixture was loaded into a gas-tight glass syringe (1750TLLX , Hamilton) and then mounted onto the 3D bioprinter as the primary printhead. A 150 µm inner diameter needle (Jensen Global) was fitted to the syringe and primed. The light-pipeline was mounted to the bioprinter as the secondary printhead, where 365nm light with 10 µW power output was emitted from the tip of a fiber-optic cannula with 0.22NA and 100 µm inner diameter. A gelatin-based non-photocrosslinkable support bath was prepared according to published methods (*3*). Briefly, 2% gelatin type B (Rousselot), 0.25% Pluronic F127 (Sigma Aldrich) and 0.5% gum Arabic (Sigma Aldrich) were consecutively added into the 50:50 (v/v) solution of distilled water at 60 ºC to 200 proof ethanol stirring at 500 RPM. Overall mixture pH was adjusted to 5.8 using 1M HCl and allowed to continue to stir overnight. Microspheres were then washed two times distilled water and stored . Before printing, 10 mL of gelatin microspheres compacted at 1000 g for 5 mins were washed twice with 10 mL of 100 mM HEPES solution including 0.5% LAP, 60 µM of tartrazine, and 25 µM methacrylated rhodamine (Methacryloxyethyl thiocarbamoyl rhodamine B, Polysciences, CAS # 669775-30-8). The overall support bath mixture was then compacted at 2000 g for 5 minutes. CAD models of the disc (where bioink is extruded) and two domains in different directions (the locations where further rastered with light following extrusion) were generated in Rhinoceros 6 software. Following the dual-head 3D bioprinting process, the slurry was washed two times with 100 mM HEPES with 1% Pen/Strep. Confocal images of the bioprinted construct were taken using Nikon A1R MP+ multiphoton confocal microscope and a 4X objective.

### Multi-Wavelength Bioprinting

GelMA support bath was prepared similar to the previous bioprinting processes with a minor change in the photofiller composition. Following the coacervation step, GelMA microspheres were resuspended within 1X PBS with 9% GelMA, 0.4 mM Ruthenium (Advanced Biomatrix, #5248) and 4 mM sodium persulfate (Advanced Biomatrix, #5248). In order to prepare NorPEG-based bioink, 20 kDa 8-arm PEG Norbornene (JenKem Chemicals) was dissolved within 1X PBS at 15% (w/v) concentration. After making sure that the chemical fully dissolved through consecutive 30 seconds sonication and 10 seconds vortexing steps, 500 µL of the solution was transferred to an Eppendorf tube. Then, 15 mM DTT and 1% LAP were added into the 15% NorPEG and then stirred for an hour in dark. The syringe with NorPEG-based bioink was mounted on the bioprinter as the primary printhead. A 150 µm inner diameter needle (30 G, Jensen Global) was attached to the tip of the syringe and then primed. The light-pipe was assigned as the secondary printhead and mounted onto the bioprinter. The tips of the primary printhead needle and fiber-optic cannula were aligned using a custom-design alignment setup. In order to provide multi-wavelength photoactivation, the light-pipeline was modified by adding a filter wheel in between collimator and fiber-optic adapter. Before each switch in between the printheads, two optical filters positioned right next to the collimator were alternated using G-code commands.

Tissue scaffolds with parallel muscle architecture were designed in Rhinoceros software and then sliced in Simplify 3D. It was sliced so for each layer, Step 1 the primary printhead extruded parallel NorPEG filaments into the slurry, Step 2 the secondary light-pipe printhead delivered light with a 405 nm long-pass filter to photocrosslink the GelMA support bath, and Step 3 the filter wheel switched to the 365 nm band-pass filter and the secondary light-pipe printhead delivered light to the extruded NorPEG filaments to photocrosslink them. This three-step process repeated in every layer until the bioprinting process was completed, and then the non-photocrosslinked support bath was melted at 37 ºC in the incubator. Constructs were then removed, washed 3 times with 1X PBS with 1% Pen/Strep, and imaged on the stereomicroscope.

**Statistical Analysis**

Statistical analysis was performed using Prism 6 (GraphPad) software, and statistical tests were chosen according to experimental conditions and data requirements. Each experiment was performed in triplicate unless otherwise specified. For statistical analysis of the half-angle (Fig. 2), evolution of effective modulus depending on printing speed and power output (Fig. 3), compaction at day 5 (Fig. 3), contractile shortening and recoiling percentage (Fig. 4) characterizations, one-way analysis of variance (ANOVA) followed by Tukey's multiple comparisons post-test was performed. Statistical significance was considered a p-value of $<0.05$.

### SUPPLEMENTAL INFORMATION REFERENCES

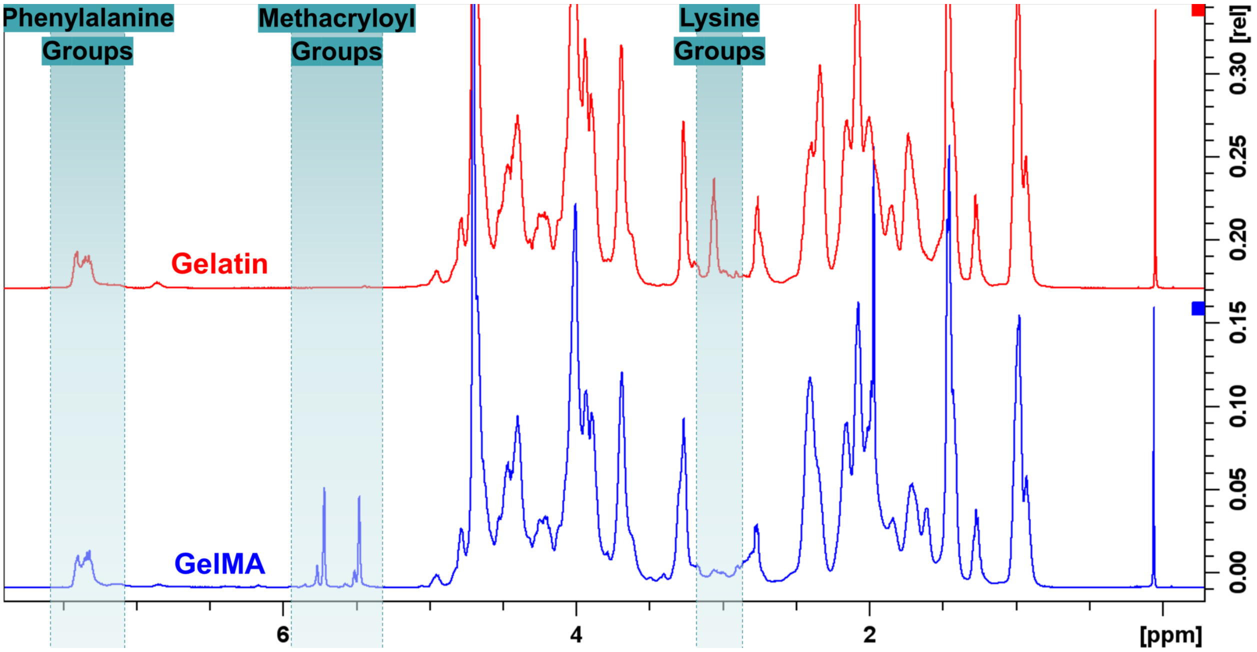


**Fig. S1. NMR characterization of the synthesized GelMA.** While phenylalanine peaks did not change during the GelMA synthesis process, the lysine peak decreased and methacryloyl peaks formed as compared to the unmodified gelatin.

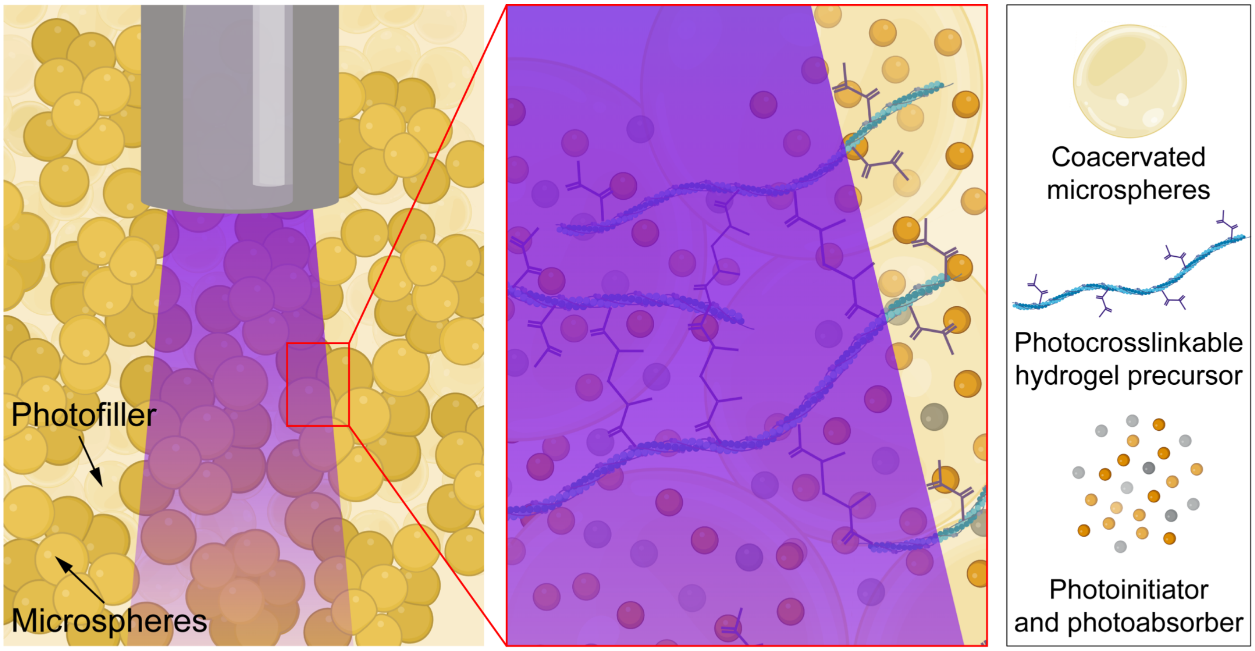


**Fig. S2. Composition of photocrosslinkable support bath.** The support bath is made out of GelMA-based microspheres and a photofiller solution including GelMA, photoinitiator and photoabsorber. During the bioprinting process, the light initiates photocrosslinking of the support bath in specific domains.

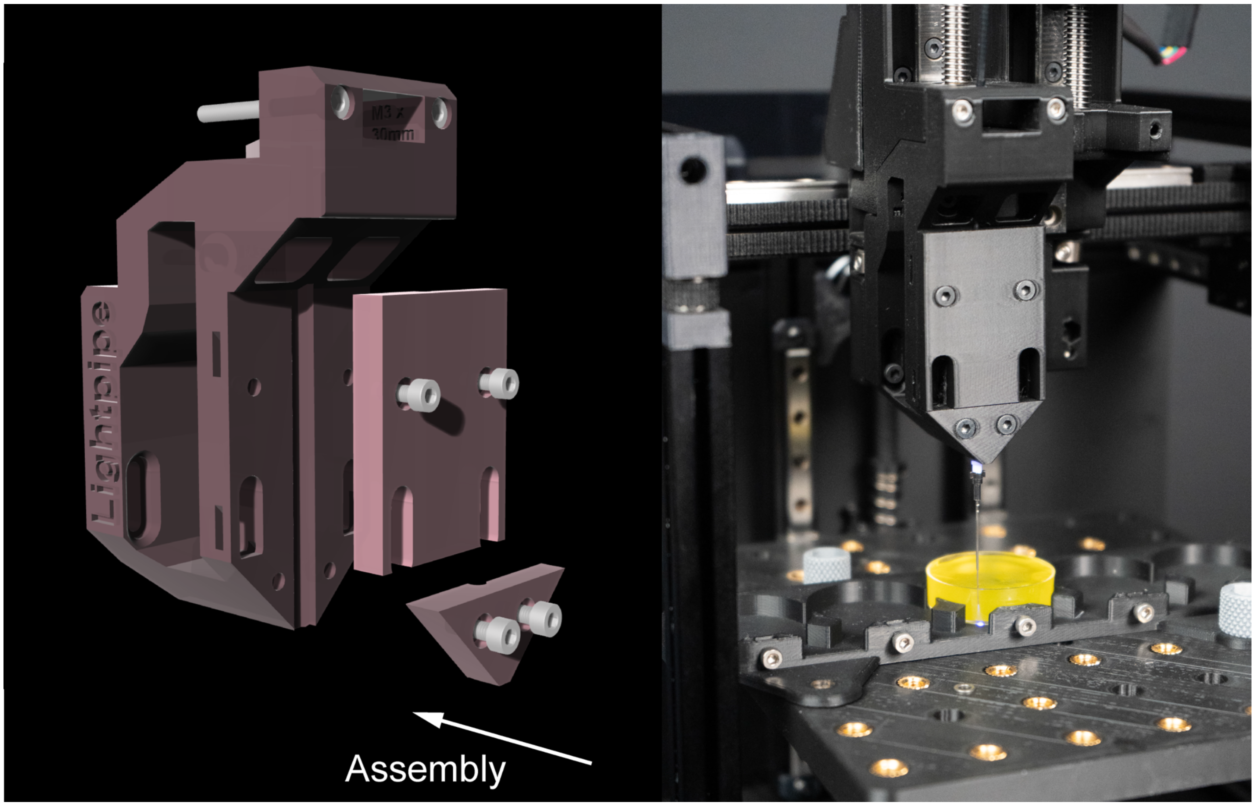


**Fig. S3. Schematic and photo of the printhead designed for integration of the light-pipeline into the 3D bioprinter.** (left) Design of the mounting hardware, where the optical fiber-coupled light path consisting of the patch cord, mating sleeve and fiber-optic cannula are fixed in place. (right) Photograph of the light-pipe setup on the 3D bioprinter.

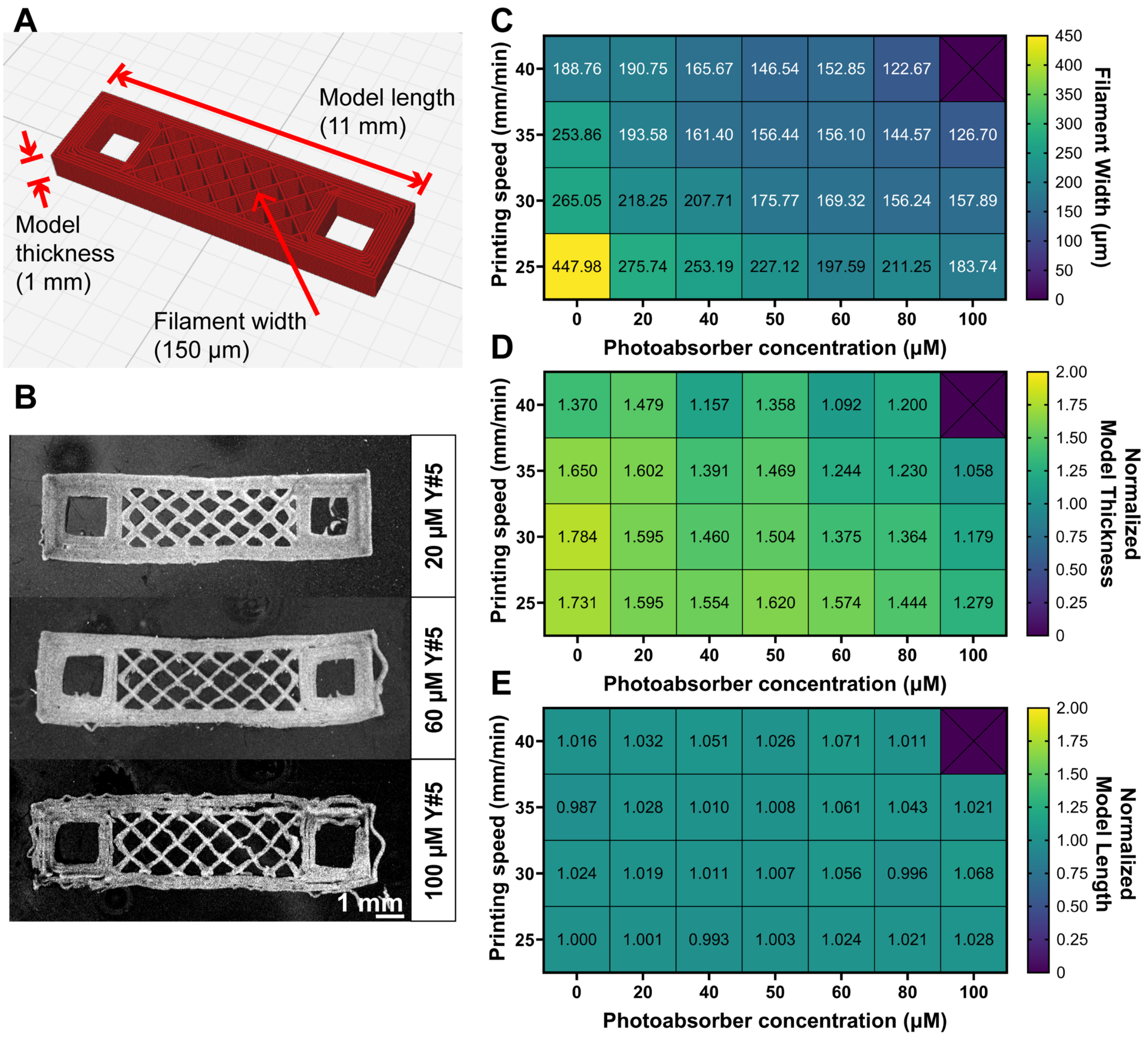


**Fig. S4. Optimization of the photoFRESH printing parameters for the photocrosslinkable support bath.** (A) Schematic of the scaffold design and of measurement locations for parameter optimization. (B) OCT images showing examples for different photoabsorber concentrations at constant print speed. (C) Filament width as a function of print speed and photoabsorber concentration, where the target dimension was 150 µm. (D) Model thickness as a function of print speed and photoabsorber concentration, where the target dimension was 1 mm. (E) Model length as a function of print speed and photoabsorber concentration, where the target dimension was 11 mm.

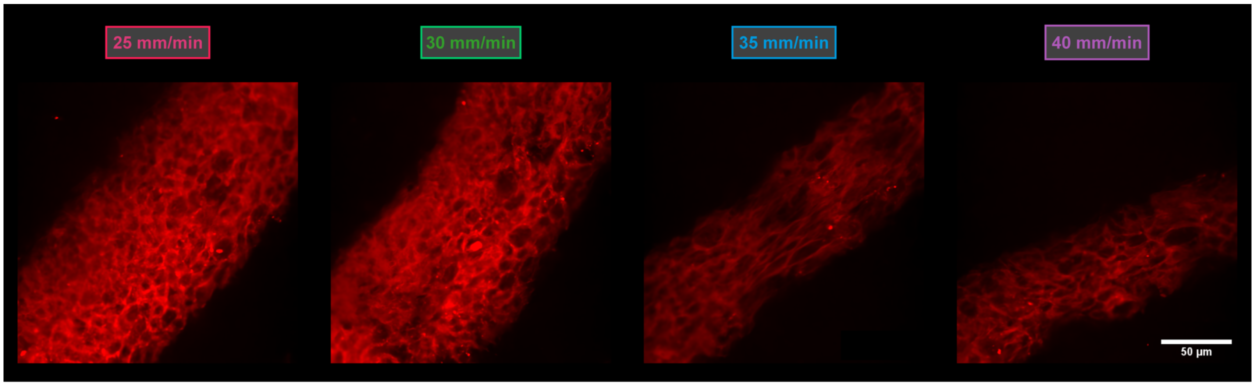


**Fig. S5. Surface morphology of filaments printed at constant light intensity different speeds.** The microindentation sample was photoFRESH printed with four domains at different printing speeds and the autofluorescence of the GelMA was imaged under confocal microscopy. The filaments had a dimple-like surface morphology across all different printing speeds. This appearance was due to the microspheres in the support bath, which leave dimple-like voids when the non-photocrosslinked GelMA support bath was melted following the photoFRESH printing process.

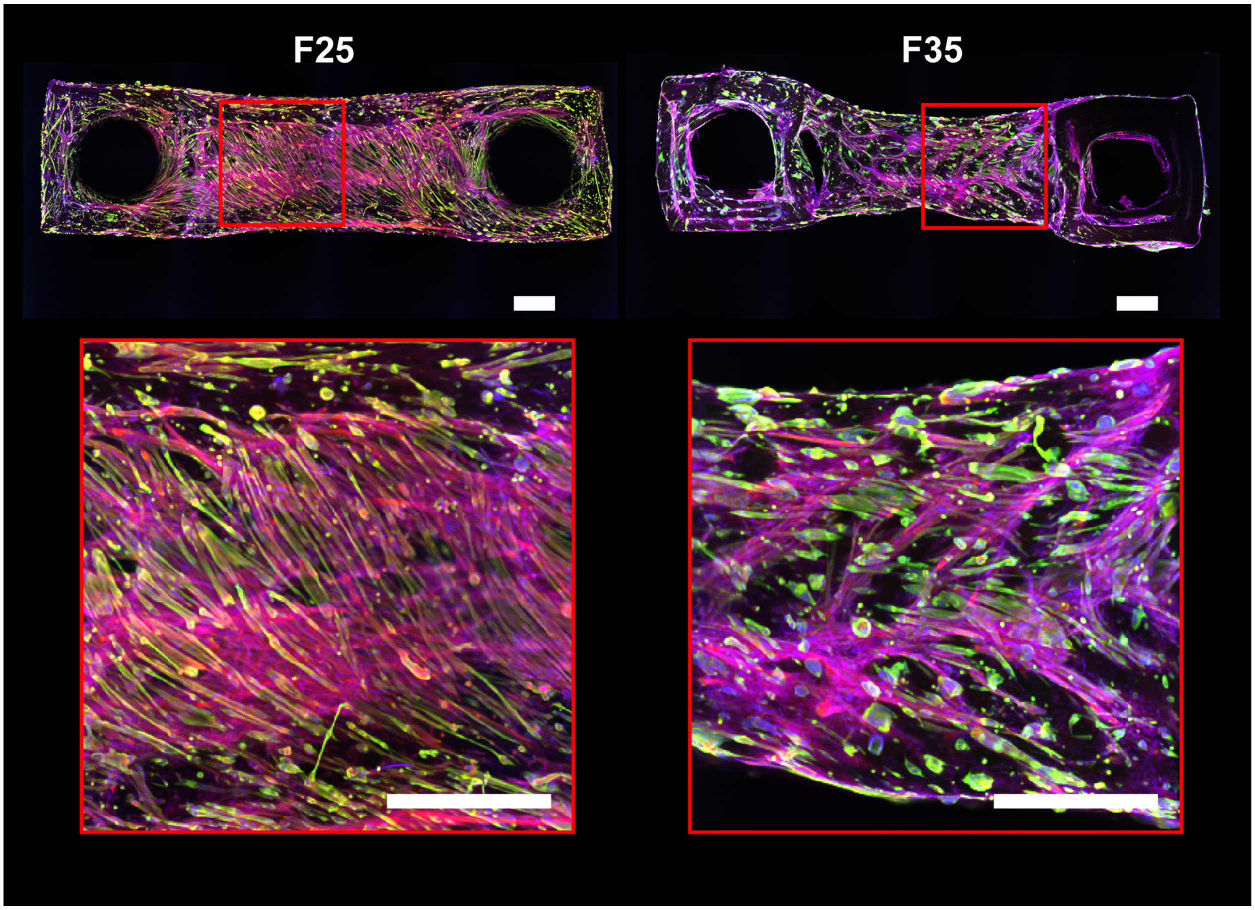


**Fig. S6. Confocal images of tissue scaffolds demonstrating the print speed dependent changes in the scaffold morphology.** Higher magnification confocal images of the tissue constructs showing the difference in C2C12 myoblast interaction with the photoFRESH printed GelMA scaffold as a function of stiff (F25) and soft (F35) printing speeds [blue: nuclei, green: myosin heavy chain, red: F-actin] (scale bars: 1 mm).

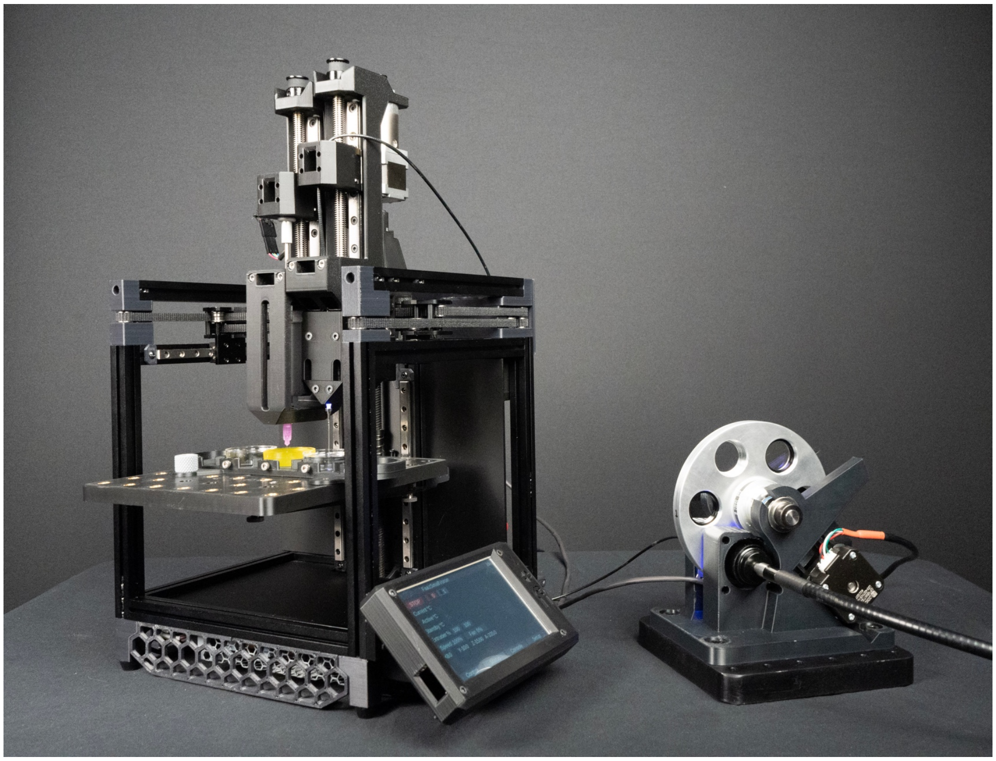

**Fig. S7. Multi-wavelength bioprinting setup where a filter wheel was incorporated into the light pipeline.** The filter wheel rotates during the photoFRESH printing process every time the required wavelength of light changes. Each position in the filter wheel can hold a different bandpass or longpass optical filter, depending on the specific photochemistry being used and the emission spectra of the connected light source.

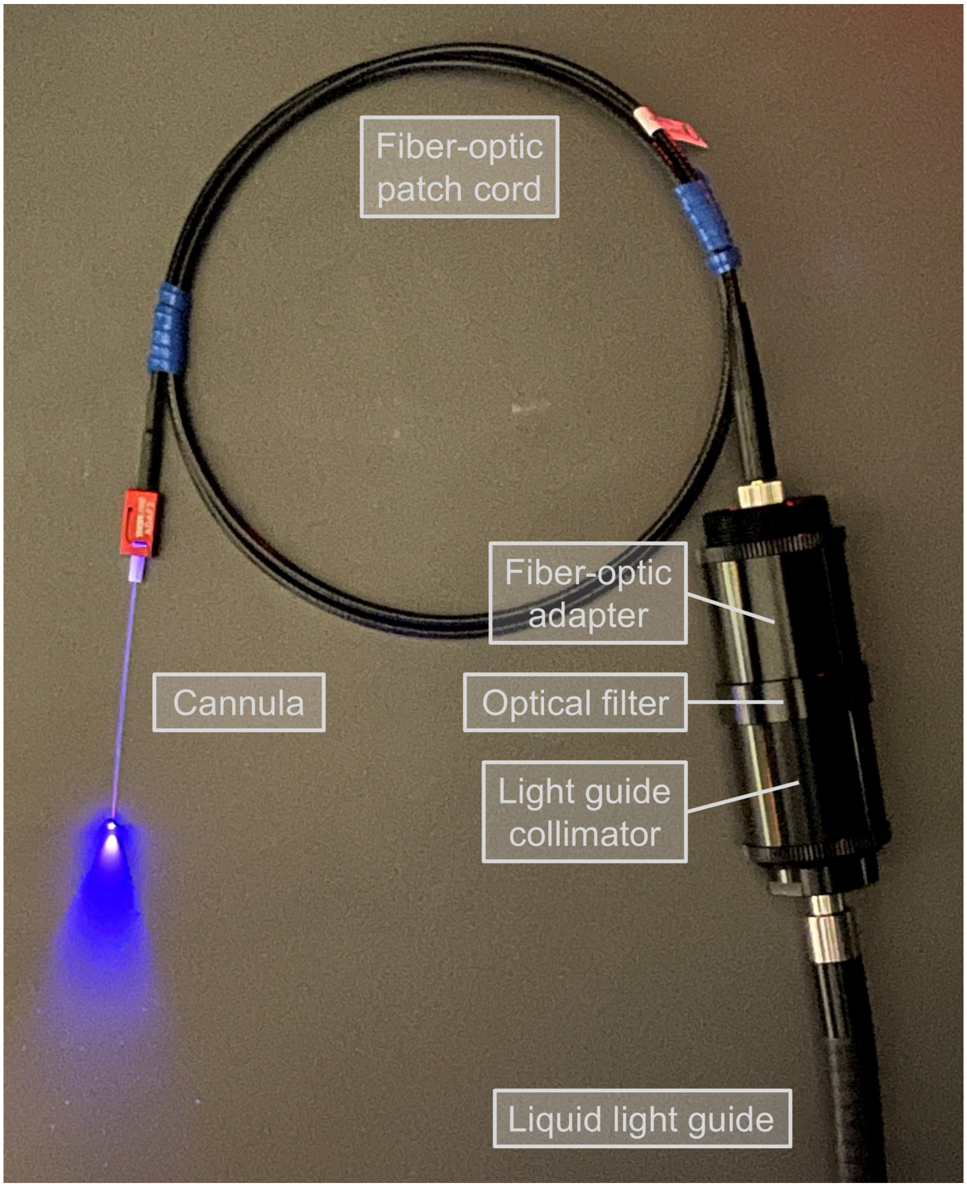


**Fig. S8. Key components of the fiber optic-coupled light path.** The UV/visible light source was connected to a collimator via a liquid light guide. The collimated light was then connected through an optical filter to pass only the specific wavelengths of light required for photoFRESH. The light was then focused into a fiber optic path cord by a fiber optic adapter. The patch cord was mated with the ceramic ferrule to the fiber-optic cannula by a mating sleeve.

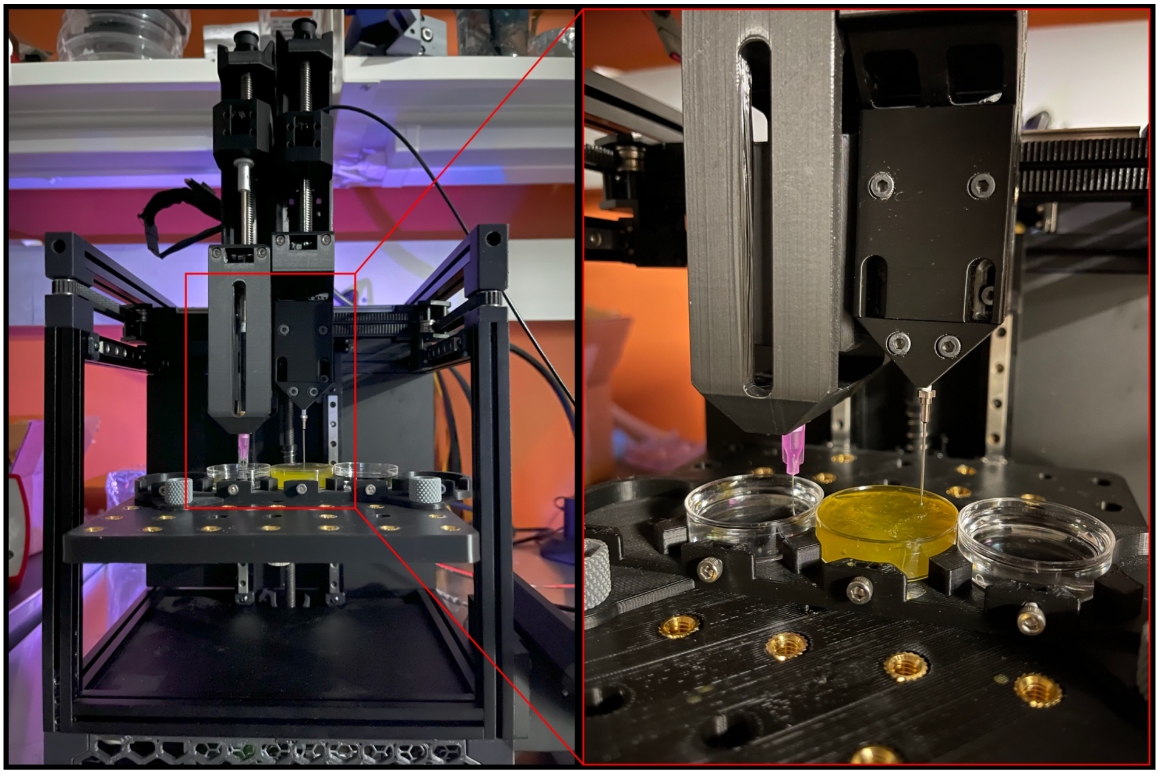

**Fig. S9. The dual-head 3D bioprinting platform combining FRESH and photoFRESH.** Overall view (left) and closer view (right) of the 3D bioprinter setup integrated with an extrusion printhead with a syringe and a light-pipe printhead with a fiber-optic cannula.

### CAPTIONS FOR MOVIES S1 TO S7

**Movie S1. PhotoFRESH 3D bioprinting in action.** Light delivered from the tip of the fiber-optic cannula is rastered within the photocrosslinkable support bath in a layer-by-layer fashion.

**Movie S2. OCT videos of overcured and optimally printed tissue scaffolds.** Spreading and scattering of light within the photocrosslinkable support bath can lead to overcuring of the previously patterned bottom layers. Optimization of the printing conditions reduce overcuring.

**Movie S3. Calcium imaging of the bipennate tissue construct printed at 25 mm/min under field stimulation.** Calcium cycling is observed under spontaneous and paced contractions of 1, 5, 10, and 20 Hz.

**Movie S4. Calcium imaging of the bipennate tissue construct printed at 35 mm/min under field stimulation.** Calcium cycling is observed under spontaneous and paced contractions of 1, 5, 10, and 20 Hz.

**Movie S5. Contraction of the tissue construct under 1Hz field stimulation after removing one of the posts.** The stiffer scaffold domains contract the less and the softer domains contract more.

**Movie S6. Top-down OCT video of myotendinous tissue-like construct fabricated through combined FRESH and photoFRESH bioprinting.** The multi-pennate architecture of the scaffold consisted of the muscle-like domain printed from the GelMA support bath by photoFRESH and the tendon-like domain printed from the collagen type I bioink by FRESH.

**Movie S7. Combined multi-modal photoFRESH and FRESH 3D bioprinting in action.** Following the extrusion of methacrylated collagen bioink into the support bath using FRESH printing, methacrylated rhodamine within the support bath was covalently coupled to the extruded collagen only in the areas exposed to light using photoFRESH.
